# Mother-to-embryo vitellogenin transport in a viviparous teleost *Xenotoca eiseni*

**DOI:** 10.1101/708529

**Authors:** Atsuo Iida, Hiroyuki Arai, Yumiko Someya, Mayu Inokuchi, Takeshi A. Onuma, Hayato Yokoi, Tohru Suzuki, Kaori Sano

## Abstract

Vitellogenin (Vtg), a yolk nutrient protein that is synthesized in the livers of female animals, and subsequently carried into the ovary, contributes to vitellogenesis in oviparous animals. Thus, Vtg levels are elevated during oogenesis. In contrast, Vtg have been genetically lost in viviparous mammals, thus the yolk protein is not involved in their oogenesis and embryonic development. In this study, we identified Vtg protein in the livers of females during the gestation of the viviparous teleost, *Xenotoca eiseni*. Although vitellogenesis is arrested during gestation, biochemical assays revealed that Vtg protein was present in ovarian tissues and lumen fluid. The Vtg protein was also detected in the trophotaenia of the intraovarian embryo. Immunoelectron microscopy revealed that Vtg protein is absorbed into intracellular vesicles in the epithelial cells of the trophotaenia. Furthermore, extraneous Vtg protein injected into the abdominal cavity of a pregnant female was subsequently detected in the trophotaenia of the intraovarian embryo. Our data suggest that the yolk protein is one of the matrotrophic factors supplied from the mother to the intraovarian embryo during gestation in *X. eiseni*. To our knowledge, this is the first report of the experimental verification of mother-to-embryo substance transport in a viviparous teleost.

## Introduction

Viviparity is a form of reproductive system in which fertilization and embryonic development progress inside the mother’s body prior to the birth, as opposed to oviparity where such developmental events are completed outside of the parent. In some viviparous vertebrates, the embryo receives nutrients from the mother in addition to those in the yolk sac. The nutrient supply and embryonic growth in the mother’s body are important factors influencing survival in the habitat environment after the birth (1). In eutherian mammals, maternal nutrients including blood plasma pass into the fetus via a placenta and umbilical cords (2). Viviparous species also occur in reptiles, amphibians, and fish (3–5). The embryos of some amphibians and sharks eat each other and develop within the mother’s body (6,7). A viviparous reptile possesses a placentalike structure similar to those found in eutherians (8). In some viviparous chondrichthyans and teleosts, a nutrient-rich liquid is supplied to the embryo in the ovary during gestation (9, 10). Diverse mechanisms of mother-to-embryo nutrient supply occur among viviparous vertebrates. However, the nature of the maternal nutrients in viviparous species is still unknown except in the eutherian mammals.

The teleost family Goodeidae is currently known to include 42 viviparous species distributed in the lakes and rivers of North America (11). Their embryo increases in weight in the mother’s body during the gestation, suggesting that maternal nutrients are supplied in addition to those in the yolk sac (5). In the mid to late gestation stages, the goodeid embryo possesses a pseudoplacenta, called the trophotaenia, around its anus. The pseudoplacenta partially consists of the intestine and is thought to be involved in nutrient incorporation during gestation. The trophotaenia consists of absorptive epithelium, blood vessels, and mesenchymal connective tissues (12, 13). Previous studies hypothesized that the ovarian fluid components are absorbed into the epithelial cells and then carried into the capillary (12, 14, 15). In *Xenotoca eiseni*, the viviparous goodeid species used in this study, the intraovarian embryo develops for approximately five weeks before birth. The embryo’s growth depends on matrotrophic nutrition absorbed from the trophotaenia for most of its intraovarian development (16–18). The trophotaenia is regressed by apoptosis during the latest stage of gestation (19); it is no longer required in the postnatal growth phase when oral food intake occurs. The components of the maternal nutrients absorbed from the trophotaenia have not been identified but are hypothesized to be secreted blood serum proteins.

Vitellogenin (Vtg) is a glycolipophosphoprotein typically present in females, but also in minor amounts in males, that is conserved in nearly all oviparous species including fish, amphibians, reptiles, birds, monotremes, and most invertebrates (20–22). In these vertebrates, *vtg* genes are typically expressed and synthesized in female liver, then the protein product is transported to the ovary via the bloodstream. In the ovary, Vtg protein is cleaved into subdomains and absorbed into the egg yolk (23, 24). In eutherian mammals, *vtg* genes have been genetically lost through co-evolution with casein genes, thus few yolk nutrients are included in their eggs (21, 25). In contrast, some viviparous fish species have maintained *vtg* genes, and a large amount of yolk nutrient is contained in their eggs (26, 27). In these fish, Vtg protein is a potential candidate for one of the maternal nutrients supplied into the intraovarian embryo. However, there is no definitive evidence that maternal Vtg protein is transported into the intraovarian embryo. In this study, we investigated Vtg transport from mother to embryo in the goodeid species, *X. eiseni*, and our results suggest that the yolk protein lost in eutherian mammals is one of the matrotrophic factors supplied to the intraovarian embryo during gestation in the viviparous teleost.

## Results

### Production and distribution of vitellogenin proteins during gestation

At the onset of gestation in goodeid species including *X. eiseni*, the main role of the ovary switches from egg production to maintenance of the embryo. During gestation, oogenesis is arrested at pre-vitellogenesis stages, thus there are no mature eggs in the ovarian lumen (Fig. 1A) and the ovary is dedicated to raising the embryo via nutrient supply. To investigate expression of vitellogenin genes (*vtgA, vtgB*, and *vtgC*) in *X. eiseni*, reverse transcription polymerase chain reaction (RT-PCR) analysis was performed using cDNA purified from the liver and gonad. In vitellogenic females, but not in male fish, *vtgA* and *B* were strongly expressed in the liver. In contrast, there were no detectable signals in testis and ovary. Expression of *vtgC* was minimal in all tissues examined in this study (Fig. S1A). Expression of the *vtg* genes in the liver was not decreased in pregnant females, despite the cessation of vitellogenesis (Fig. 1B). In the ovary, *vtg* genes were not expressed during gestation, as for the vitellogenesis stage (Fig. 1B, see also Fig S1A). VtgA and B proteins were detectable by immunohistochemistry in the non-pregnant female liver (Fig. S1B). The strong signals for Vtg proteins in female liver were replicated using another antibody, anti-Vtg#1 (Fig. S2A, see also Table S1). In contrast, specific signals for Vtg protein in male liver were weaker than those in female liver (Fig. S1C). The strong signals for VtgA and B proteins were also observed in the pregnant female liver (Fig. 1C). Furthermore, the Vtg proteins were detectable in the ovarian septum of the pregnant female (Fig. 1D, see also Fig. S2B). In vitellogenic females, the Vtg proteins were detectable in the ovarian epithelium approximal to the developing oocytes (Fig. S1D, see also Fig. S2C). Thus, the signals observed using the antibodies indicated endogenous Vtg protein presence and distribution in the *X. eiseni* tissues. These results prompt the question: why are the Vtg proteins present in the ovary during gestation?

**Figure 1.**
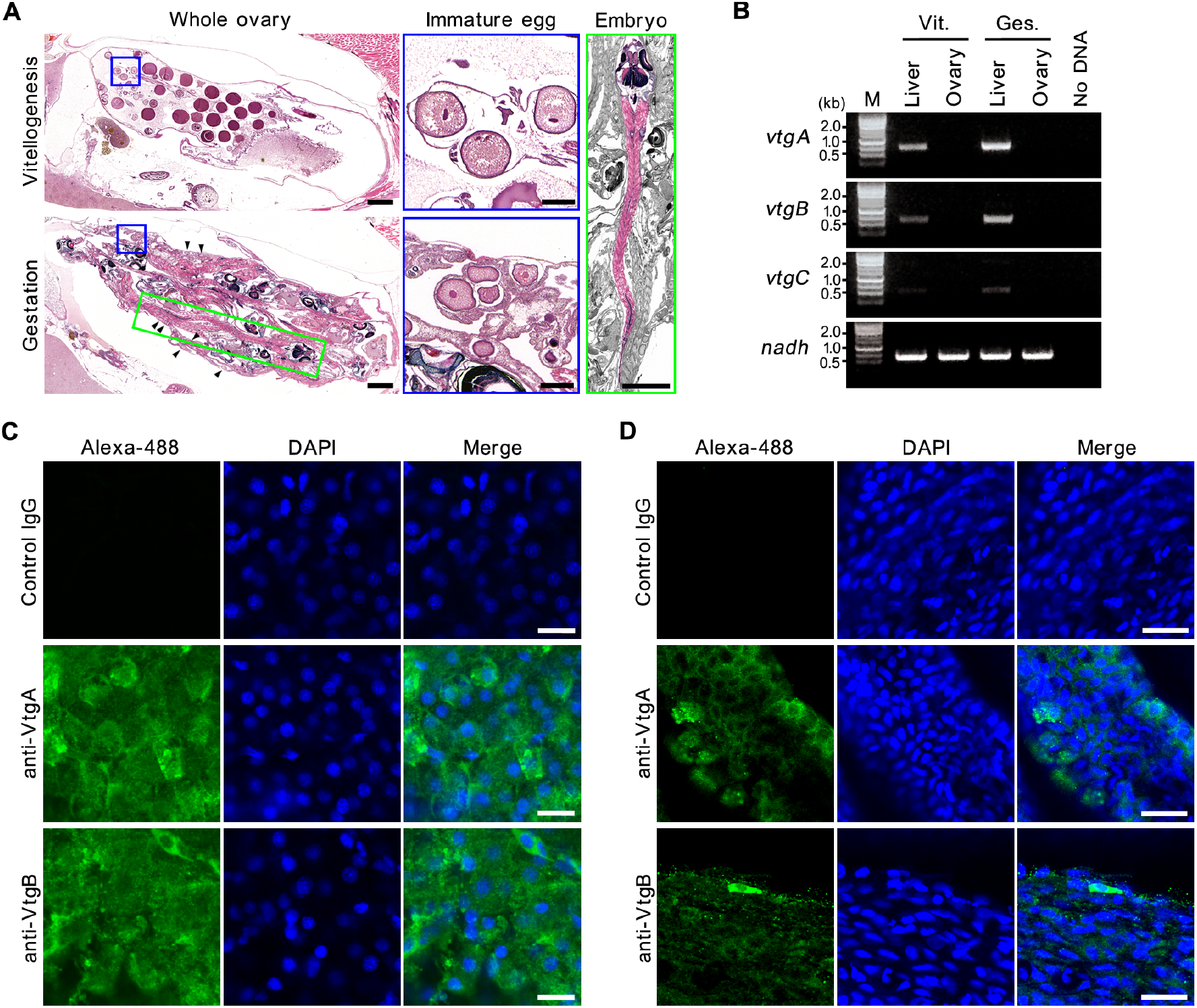
Vitellogenin synthesis and supply to ovary during gestation. **A**. HE-stained transverse sections of vitellogenesis (upper) and gestation (lower) stages of an *X. eiseni* female ovary. In the non-pregnant ovary, mature eggs including yolk were observed in the ovarian lumen, and some immature oocytes before vitellogenesis were observed on the ovarian epithelium. In the pregnant ovary, developing embryos filled the ovarian lumen, and there were no mature eggs with yolk. Arrowheads indicate immature oocytes in the ovarian epithelium. Scale bar, 1 mm (whole ovary and embryo) and 200 μm (immature egg). **B**. RT-PCR for *vtg* genes in female *X. eiseni*. All *vtg* genes were detected in the liver during vitellogenesis. The hepatic expression was also observed in the 3^rd^ week of gestation. No *vtg* expression was observed in any ovary samples. *Nicotinamide adenine dinucleotide (nadh)* was used as an internal control. M, size marker. Vit., vitellogenesis. Ges., gestation. **C**. Fluorescence immunohistochemistry for VtgA or B in lobe edges of female liver in the 3^rd^ week of gestation. Scale bar, 20 μm. **D**. Fluorescence immunohistochemistry for VtgA or B in the ovarian septum in the 3^rd^ week of gestation. Scale bar, 20 μm. See also Figures S1 and S2.

### Presence of vitellogenin proteins in intraovarian embryo

We hypothesized that the Vtg proteins in the pregnant ovary are absorbed into the intraovarian embryo as a matrotrophic factor in *X. eiseni*. To verify that, we investigated the expression of *vtg* genes and the presence of Vtg proteins in the intraovarian embryo. The *vtg* genes were not expressed in the whole embryo or the trophotaenia extracted from a pregnant female in the 3^rd^ week after mating (Fig. 2A). Thus, the intraovarian embryo would not synthesize the Vtg proteins autonomously at that stage. However, fluorescent immunohistochemistry indicated the presence of VtgA and B proteins in the epidermal layer and mesenchymal region of the trophotaenia (Fig. 2B, see also Fig. S2D). Furthermore, immunoelectron microscopy revealed that Vtg proteins are distributed in intracellular vesicles in the epithelial cells, and the mesenchymal cell surface (Fig. 2C). The signals on the mesenchymal cell surface were in proximity to an extracellular-matrix (ECM) labelled by a fibronectin antibody, but most were not overlapped (Fig. 2D). These results indicate that the Vtg proteins could be absorbed as macromolecules into the trophotaenia though the epithelial cells, retaining their antigenicity.

**Figure 2.**
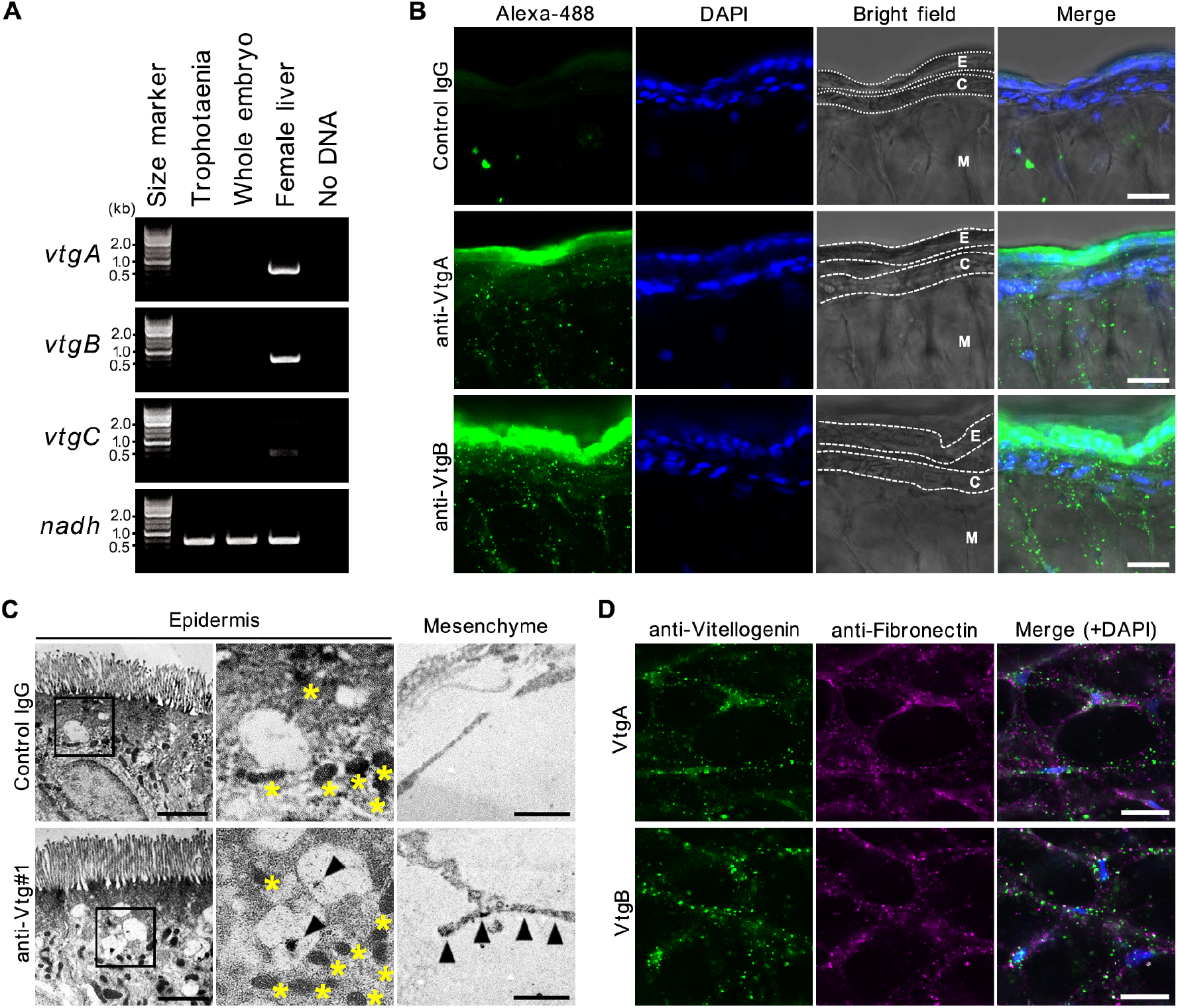
Vitellogenin distribution in the trophotaenia of the intraovarian embryo. **A**. RT-PCR for *vtg* genes in *X. eiseni* intraovarian embryo in the 3^rd^ week postfertilization. There was no *vtg* expression in whole embryo and trophotaenia. Female liver was used as a positive control for the reaction. *Nicotinamide adenine dinucleotide (nadh)* was used as an internal control. **B**. Fluorescence immunohistochemistry for VtgA or B in the trophotaenia of the intraovarian embryo in the 3^rd^ week postfertilization. E, epidermal layer. C, capillary. M, mesenchyme. Scale bar, 20 μm. **C**. Immunoelectron microscopy using anti-Vtg#1 for the trophotaenia of the intraovarian embryo in the 3^rd^ week post-fertilization. Enlarged image for epidermis show specific signals against Vtg in vesicles (arrowheads). Dark spots in the cytoplasm marked with asterisks indicate mitochondria. Enlarged image for mesenchyme shows specific signals on the mesenchymal cell (arrowheads). Scale bar, 1 μm. **D**. Dual fluorescence immunohistochemistry for Vtg and fibronectin (a marker for extracellular matrix). Scale bar, 20 μm. See also Figure S2.

### Secretion of vitellogenin proteins into ovarian fluid

The intraovarian embryo is not adhered to the maternal tissues in *X. eiseni*. Thus, the maternal components including Vtg proteins are thought to dissolve in the ovarian fluid as secreted proteins. Coomassie brilliant blue (CBB) staining displayed the secreted proteins integrated into the ovarian fluid (Fig. 3A). A 75-kDa protein was a major component in the ovarian fluid. The second major 240-kDa protein signal could be regarded as full length Vtg proteins with post-translational modifications. Some minor protein signals underwent change between vitellogenesis and gestation. To identify signals for Vtg proteins integrated into the ovarian fluid, we performed western blotting using a Vtg antibody (Fig. 3B). We obtained six signals (240-, 100-, 75-, 55-, 50-and 45-kDa) in the ovarian fluid during gestation, including two major proteins (240-and 75-kDa) displayed by CBB staining. This result was replicated using a different Vtg antibody (Fig. S3A). Thus, the six signals were candidates for Vtg proteins or cleaved fragments integrated into the ovarian fluid. Four of the six candidate signals (100-, 75-, 50-and 45-kDa) were detectable in lysate samples from the trophotaenia of the intraovarian embryo (Fig. 3B). The 75-kDa protein was also one of the major proteins detected in whole ovary lysate (Fig. S3B). Western blotting indicated that four of the six signals (240-, 100-, 75-and 50-kDa) observed in the ovarian fluid were also detected in whole ovary lysate. These signals were female specific or grossly higher than those in the male tissues (Fig. S3C). These results indicate that the 100-, 75-and 50-kDa Vtg fragments are secreted into the ovarian fluid from the ovarian tissues without any changes in their molecular weight, and then absorbed into the embryo via the trophotaenia.

**Figure 3.**
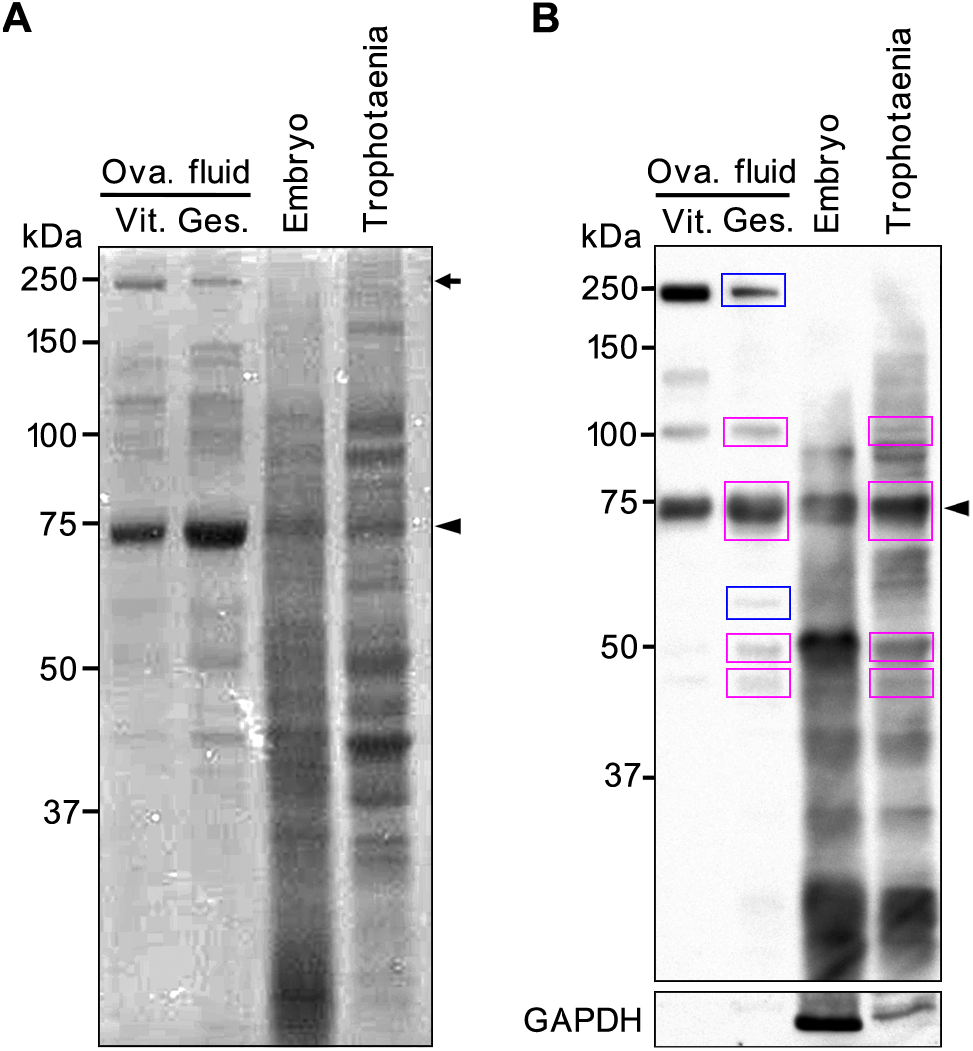
Detection of vitellogenin fragments in ovarian fluid and trophotaenia. **A**. Electrophoresis and CBB staining of ovarian fluid and intraovarian embryo lysate extracted from pregnant female fish in the 3^rd^ week after mating. The ovarian fluid showed two major protein bands of 240-and 75-kDa (arrow and arrowhead). The 75-kDa band was also observed in the lysate from whole embryo and trophotaenia. The 240-kDa protein was not detected in the embryo samples. Ova., ovarian. Vit., vitellogenesis. Ges., gestation. **B**. Western blotting (using anti-Vtg#1 antibody) of ovarian fluid and intraovarian embryo lysate extracted from pregnant female fish in the 3^rd^ week after mating. The ovarian fluid indicated two major signals against Vtg at 240-and 75-kDa (arrow and arrowhead). In the gestation sample, six specific signals (240-, 100-, 75-, 55-, 50-and 45-kDa) were observed as distinct bands. Magenta boxes indicate the signals against Vtg detected in the both ovarian fluid and trophotaenia. Blue boxes indicate the signals detected only in ovarian fluid. GAPDH was used to check for contamination of cellular components into the ovarian fluid and non-specific degradation of extracted proteins during the sample collection. See also Figure S3.

### Vitellogenin transfer from mother to intraovarian embryo

As an experimental verification the transfer of Vtg from mother to embryo, we performed a tracer assay using a fluorescence-labelled Vtg protein. We prepared a FITC-conjugated Vtg protein (FITC-Vtg) that originated from goldfish. The FITC-Vtg injected into the abdominal cavity of female fish was carried and secreted into the ovarian fluid, and then imported into the yolk (Fig. S4A, B and C). This indicated that the FITC-Vtg was functional for the mother-to-egg Vtg transfer. To validate a mother-to-embryo Vtg transfer, the tracer was injected into a pregnant female in the 3^rd^ week after mating, and then the intraovarian embryo was extracted and the trophotaenia was observed using fluorescent microscopy (Fig. 4A). The fluorescent dye injected into the abdominal cavity of pregnant female fish exuded to the vascular lumen and was then visualized in the blood vessel network including the caudal fin (Fig. 4B). Thus, both the exogenous and endogenous Vtg proteins would be transported via the bloodstream. The FITC fluorescence was accumulated in the digestive tract of the embryo (Fig. S4D). Confocal microscopy indicated that the FITC fluorescence was also detected in the epidermal layer of the trophotaenia in the embryo extracted from the FITC-Vtg injected female. In contrast, there were no signals in the PBS-injected control. Furthermore, the fluorescence was merged to immunohistochemistry signals against FITC (Fig. 4C). This means that the intraovarian embryo incorporated exogenous Vtg protein into the epidermal cells of the trophotaenia from the mother’s body without loss of the fluorescence and antigenicity of labelled FITC. We also confirmed that a FITC-conjugated dextran (M.W. 250,000) exuded to the vasculature and was absorbed into the trophotaenia (Fig. S5). These results indicated that maternal proteins and carbohydrates in interstitial fluid and blood serum could be transferred into the intraovarian embryo via the trophotaenia (Fig. 5).

**Figure 4.**
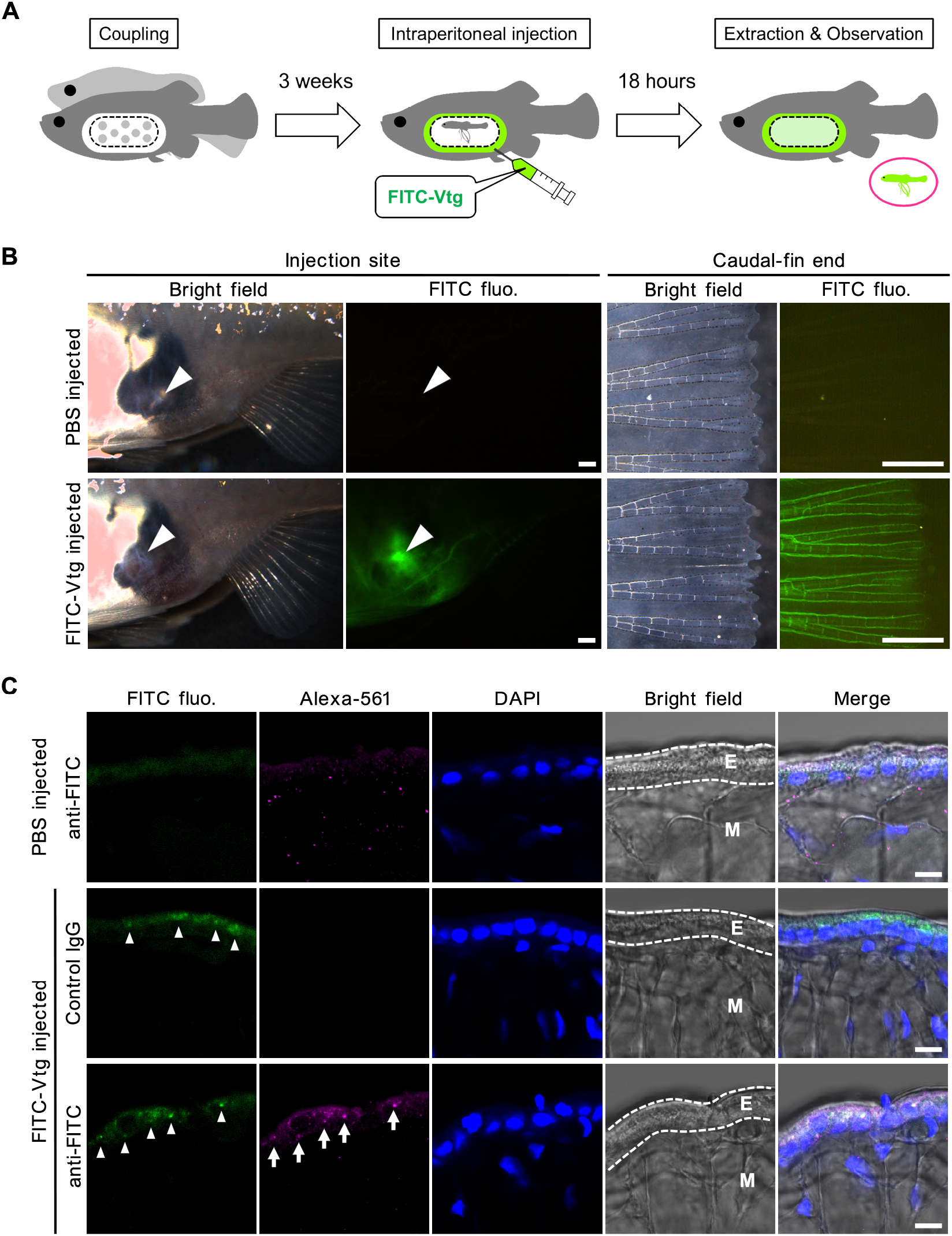
Tracing for mother-to-embryo vitellogenin transfer in live fish. **A**. A scheme for the tracing analysis. Mating: To prepare a pregnant female, a pair of mature *X. eiseni* were mated, and then the male fish was removed after mating. Intraperitoneal injection: At 3 weeks after mating, fluorescent Vtg solution was injected into the abdominal cavity of the pregnant female. Extraction & observation: At 18 hours after injection, the embryo was extracted from the female fish and observed under a fluorescent microscope. **B**. Fluorescent microscopy for the injected fluorescent solution in the female fish. The solutions were injected into the abdominal cavity from the gravid spot of pregnant female fish (arrowhead). The blood vessels in the caudal fin were visualized by a green fluorescence for the FITC-Vtg. Scale bar, 1 mm. **C**. Fluorescent microscopy of the intraovarian embryo. Green fluorescence was detected in the epidermal cell layer of trophotaenia in the extracted embryo (arrowheads). The green fluorescence was merged to signals for anti-FITC (arrow). E, epidermal layer. M, mesenchyme. Scale bar, 10 μm. See also Figures S4 and S5.

**Figure 5.**
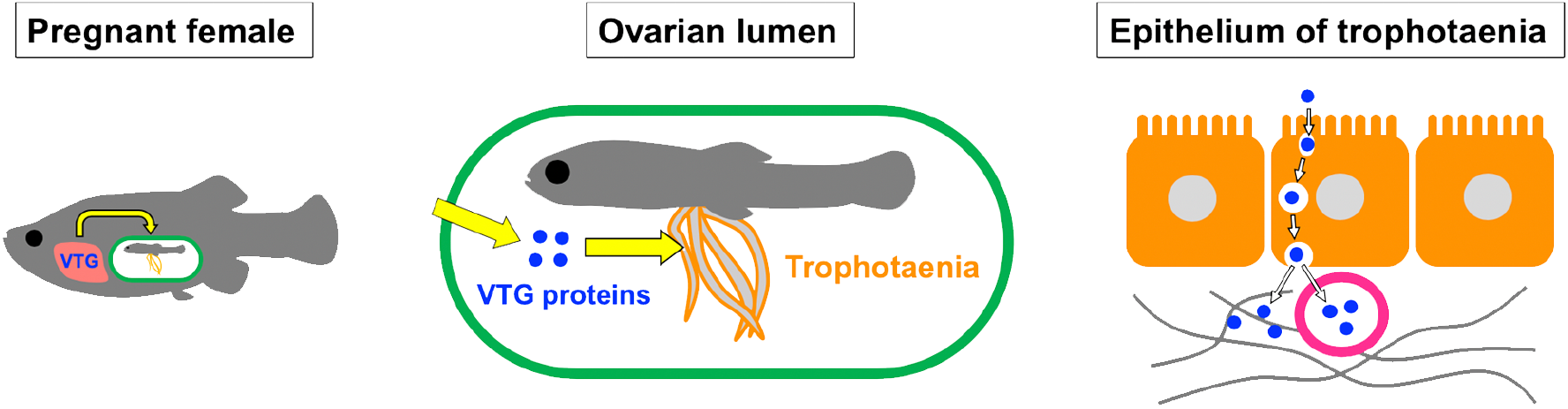
A model for mother-to-embryo vitellogenin transfer during gestation. In the pregnant female, vitellogenin genes are strongly expressed in the liver, and then the proteins are carried to the ovary via the bloodstream. In the ovarian lumen, full length and fragmented vitellogenin proteins are secreted into the ovarian fluid, and then absorbed into the embryo via the epithelium of the trophotaenia. In the epithelium of the trophotaenia, the fragmented vitellogenin proteins are imported into the epidermal cells and carried as macromolecules via vesicle trafficking.

## Discussion

In this study, we revealed Vtg synthesis and mother-to-embryo transfer during gestation in the goodeid viviparous teleost *X. eiseni* (Fig. 5). In some viviparous species, Vtg has been considered to unnecessity for gestation, and the need for and roles of Vtg in gestation were mostly elusive in viviparous teleosts. A previous study reported that Vtg-like proteins did not appear in the serum of ovarian fluid during the gestation period of viviparous surfperches (28). Another study indicated that Vtg content in female serum is downregulated during gestation in a viviparous eelpout (29). The eelpout exhibits a higher Vtg level during vitellogenesis than that during gestation. This suggests that Vtg is required for oogenesis, and that supply into the oocytes could be regulated according to the reproduction cycle. In the family Goodeidae, a previous study indicated that Vtg is not a maternal nutrient supplied to the intraovarian embryo throughout gestation in *Goodea atripinnis* and *Alloophorus robustus* (30). In contrast, other research group argued that Vtg protein was utilized for intraovarian embryonic growth in *Girardinichthys viviparus* and *Ameca splendens*, however, definitive evidence of mother-to-embryo transfer of Vtg protein has not been demonstrated (27). In the present study, we revealed that *Vtg* gene expression and protein secretion are maintained at a high level during vitellogenesis and gestation in *X. eiseni*. Furthermore, the tracer analysis revealed that FITC-conjugated Vtg protein could be transferred into the trophotaenia of the intraovarian embryo from the mother. The accumulation of the FITC-fluorescence in the digestive tract might indicated a destination of the Vtg protein or its degradants absorbed from the trophotaenia. This evidence suggests that Vtg is one of the matrotrophic factors in *X. eiseni*.

We performed the western blotting analysis to identify the Vtg proteins in the ovarian fluid and the intraovarian embryo by their molecular weight. The molecular weights of the intact VtgA and B proteins in *X. eiseni* were predicted to be approximately 190 kDa. Following synthesis, Vtg proteins undergo post-translational modifications like glycosylation and phosphorylation, and subsequently lipidation (31). Thus, the actual weight of native Vtg proteins is difficult to predict. The CBB staining and western blotting indicated that the 75-kDa Vtg fragment is the major protein secreted into the ovarian fluid, and our data revealed that the 100-, 75-and 50-kDa Vtg fragments were absorbed into the trophotaenia. Previous studies indicated that Vtg protein undergoes a specific proteolytic cleavage during uptake into an oocyte (23). Vertebrate Vtg protein is cleaved into the heavy (~120-kDa) and light (~45-kDa) chain lipovitellin, and phosvitin (~34-kDa) (24). We used polyclonal antibodies against whole Vtg sequence purified from blood plasma of 17β-estradiol treated arctic char or sea bream. Thus, we could not identify a correspondence relationship between the signals and predicted domain types. Additionally, some smaller bands and smear-like signals against the antibodies were observed in the embryo and trophotaenia. A previous study indicated that Vtg protein in zebrafish embryo was digested and expended during development (32). Therefore, we suggest that the lower signals are Vtg fragments digested to be expended as a maternal nutrient.

Immunoelectron microscopy revealed that trophotaenia epidermal cells import Vtg fragments via vesicle trafficking. Similar protein uptake by lysosomes or endosomes was previously detected by absorption assay of cationized ferritin (CF) or horseradish peroxidase (HRP) (14, 15). We also demonstrated that high-molecular weight dextran is absorbed into the trophotaenia. These findings suggest that the epidermal layer of the trophotaenia functions as an absorptive epithelium that imports maternal proteins and other components as macromolecules. The trophotaenia is partially derived from the hindgut and exhibits structural and functional characteristics of intestinal absorptive cells (12, 13). Similar endocytotic absorption of a macromolecule was previously reported in suckling rat absorptive cells (33–35). The suckling stage-specific absorption undergoes changes corresponding with the development of the digestive system (36). Therefore, we hypothesize that the macromolecule absorption observed in viviparous fish species is also a common characteristic in the post-embryonic stage of vertebrates. Mammals use this mechanism to assimilate macromolecules from the mother’s milk, whereas in goodeid fish the trophotaenia, which is a modified juvenile hindgut, is utilized for uptake of the intraovarian macromolecules supplied from the mother. To validate this hypothesis, investigation and comparison of the molecular mechanisms of the absorption system in juvenile stages of mammals and viviparous teleosts is required, and furthermore in other species.

## Acknowledgements

Breeding and experiments were performed in the laboratory of Dr. Atsuko Sehara-Fujisawa. We thank Yoshiko Aihara for helpful discussion. Ryo Kurokawa helped with the sample collection. This work was supported by crowdfunding via The Academist, Inc. The crowdfunding investors for this project were Nobuo Ishii, Kyoko Kawamura, Mayu Miyamoto, Eiri Ono, Shoko Saito, Kohei Shibata, Tomosato Takabe, Yuma Nihata, Tokuyoshi Wakamatsu, Yoshinori Wakamatsu, Shoichi Tamura, Keiko Grace Kobori, Tsuneaki Hasekura, and the Patchwork Club of Fujimidai Elementary School (Nagoya, Japan).

## Author contributions

A.I. designed and performed the experiments. K.S. helped with the experimental design and handled the genetic cloning. Y. S., M.I., and T.A.O. contributed to a part of the experiment. H.Y. and T.S. prepared the fluorescent vitellogenin protein. A.I. wrote the manuscript, and all authors approved the final version submitted.

## Methods

### Animal experiments

This study was approved by the ethics review board for animal experiments of Kyoto University. We sacrificed live animals in minimal numbers under anesthesia according to the institutional guidelines.

### Fish breeding

*Xenotoca eiseni* was purchased from Meito Suien Co., Ltd. (Nagoya, Japan). Adult fish were maintained in freshwater at 27°C under a 14:10-h light: dark photoperiod cycle. Fish were bred in a mass-mating design, and approximately 50 fish were maintained for this study. The juveniles were fed live brine shrimp larvae and Hikari Rabo 450 fish food (Kyorin Co., Ltd., Himeji, Japan), and the adults were fed Hikari Crest Micro Pellets (Kyorin). To accurately track the pregnancy period, the laboratory-born fish were crossed in a pair-mating design, and the mating behavior was recorded.

### Sample collection

Fish samples were anesthetized using tricaine on ice, prior to surgical extraction of tissues or embryos. The obtained samples were stored on ice until the subsequent manipulations. In this study, we dissected approximately 20 adult fish and 30 pregnant females, and extracted 15–30 embryos in each operation.

### Antibodies

Polyclonal antibodies against VtgA and B were generated in this study. The antigen sequences are aa 901-1,018 (VtgA, GenBank: ACI30217.1) and aa 902-993 (VtgB, GenBank: ACI30218.1). Inclusion bodies of the antigen peptides were harvested from the transformant *Escherichia coli* BL21 Star (DE3) (Thermo Fisher Scientific, Waltham, MA, USA) (37). For determination of N-terminal sequences of the antigen protein, the inclusion bodies loaded onto Ni-NTA Superflow (Qiagen, Valencia, CA, USA) were subjected to a protein sequencer (Procise 491HT; Applied Biosystems, Foster City, CA, USA). 0.2mg of each inclusion body diluted in urea/PBS was injected into Jcl:ICR mice (CLEA Japan, Inc., Meguro, Japan) four times every two weeks. Serum samples were harvested from the antigen-injected mice. The serums, diluted to 50% in glycerol with 0.1% sodium azide, were used for immunohistochemistry. Information for the commercial antibodies used is listed in Table S1.

### Reverse transcription (RT) PCR

Total RNA was extracted from tissues or whole embryo using the RNeasy Mini kit (Qiagen) and reverse-transcribed using SuperScript III reverse transcriptase (Thermo Fisher Scientific). PCR was carried out using KOD-FX (Toyobo, Osaka, Japan) under the following conditions: 100 s at 94°C, followed by 28 cycles of 20 s at 94°C, 20 s at 60°C, and 20 s at 72°C; and 40 s at 72°C. Primer sequences are listed in Table S1.

### Fluorescent Immunohistochemistry

Tissue samples were fixed in 4.0% paraformaldehyde/phosphate-buffered saline (PFA/PBS) at 4°C overnight. When using the anti-Vtg#1, fixed samples were incubated in 10 mM Tris-EDTA (pH 9.5) at 95°C for 20 min as an antigen activation. The activation is not required for the other antibodies. Samples were permeabilized using 0.5% TritonX-100/PBS at room temperature for 30 min, and then treated with Blocking-One solution (Nacalai Tesque, Kyoto, Japan) at room temperature for 1 h. Primary antibodies were used at 1:100 (anti-FITC) or 1:500 (the others) dilution with Blocking-One solution. Samples were reacted with primary antibodies at 4°C overnight. Secondary antibodies (see Table S1) were used at 1:500 dilution in 0.1% Tween-20/PBS with DAPI (Sigma-Aldrich, St. Louis, MO, USA). Samples were treated in the secondary antibody solution at 4°C overnight. Microscopic observation was performed using Leica TCS SP8 microscopes (Leica Microsystems, Wetzlar, Germany).

### Immunoelectron microscopy

Embryo samples were fixed in 4.0% PFA/PBS. Fixed samples were washed in PBS and then incubated in 10 mM Tris-EDTA (pH 9.5) at 95°C for 20 min as an antigen activation. Activated samples were reacted with primary antibody (anti-Vtg#1) at 4°C overnight, and then reacted with biotinylated anti-rabbit IgG (Vector, Burlingame, CA, USA) at room temperature for 2 h. Samples were performed with the avidin–biotin–peroxidase complex kit (Vector), and visualized with 0.05% 3,3’-diaminobenzidine (DAB; Dojindo Laboratories, Kumamoto, Japan) and 0.01% hydrogen peroxide in 50 mM Tris buffer (pH 7.2) at room temperature for 10 min. The stained samples were fixed in 1% osmium tetroxide in 0.1 M sodium phosphate buffer (pH 7.4) at room temperature for 1 h. The fixed samples were dehydrated in ethanol, transferred to propylene oxide, and embedded in Spurr’s resin (Polysciences, Warrington, PA, USA). Ultrathin sections were cut with a diamond knife every 70 nm and mounted on grids. The sections were stained with uranyl acetate and lead citrate, and examined with a transmission electron microscope (JEM2100, JEOL, Tokyo, Japan).

### Western blotting

Samples were washed with cold PBS and lysed in RIPA Buffer (10 mM Tris-HCl, pH 7.4, 150 mM NaCl, 1 mM EDTA, 0.1% SDS, 1% NP-40, 0.1% sodium deoxycholate) containing 1% protease inhibitor cocktail (Nacalai Tesque). The lysates were diluted by 50% in 2 x sodium dodecyl sulphate buffer (4% SDS, 20% glycerol, 0.002% bromophenol blue, 125 mM Tris-HCl, pH 6.8) containing 10% 2-mercaptoethanol (2-ME), and denatured at 95°C for 5 min. The sample solutions were loaded and separated on 10% polyacrylamide gels, after which the proteins were transferred to polyvinylidene difluoride membranes (Immobilon-P; Millipore, Billerica, MA, USA). After blocking with Blocking-One solution (Nacalai Tesque), the membranes were incubated at 4°C overnight with primary antibodies (Vtg#1, 1:2,000, Vtg#2, 1:2,000, anti-GAPDH, 1:2,000). After washing, membranes were reacted with secondary antibody (anti-rabbit IgG, 1:5,000, Cell Signaling Technology, Danvers, MA). The signals were visualized using ECL Prime Western Blotting Detection Reagent (GE Healthcare, Chicago, IL, USA). Image data were acquired with an ImageQuant LAS 4000 (GE Healthcare).

### Tracing of mother-to-embryo transport

A FITC-conjugated goldfish Vtg protein was prepared according to the previous study (38). The FITC-Vtg, FITC-dextran (MW 250,000, Sigma-Aldrich) diluted in PBS (final conc. 1.0 mg/ml), or control solvent (PBS) was injected into the abdominal cavity of adult females under anesthesia. The injected females were incubated in a separate tank. For angiography of the injected dye, the female fish were observed under anesthesia using a Leica M205C microscope at 30 min post-injection. To investigate absorption into the ovary or intraovarian embryo, the sample was surgically harvested from the females under anesthesia at 18-, 42-or 90-hours post-injection. The extracted samples were observed using a Leica M205C, MZ16FA, or TCS-SP8 microscope.

## Supplemental Information

The supplementary information includes five Additional Figures and one Table.

### Additional Figures

**Figure S1.**
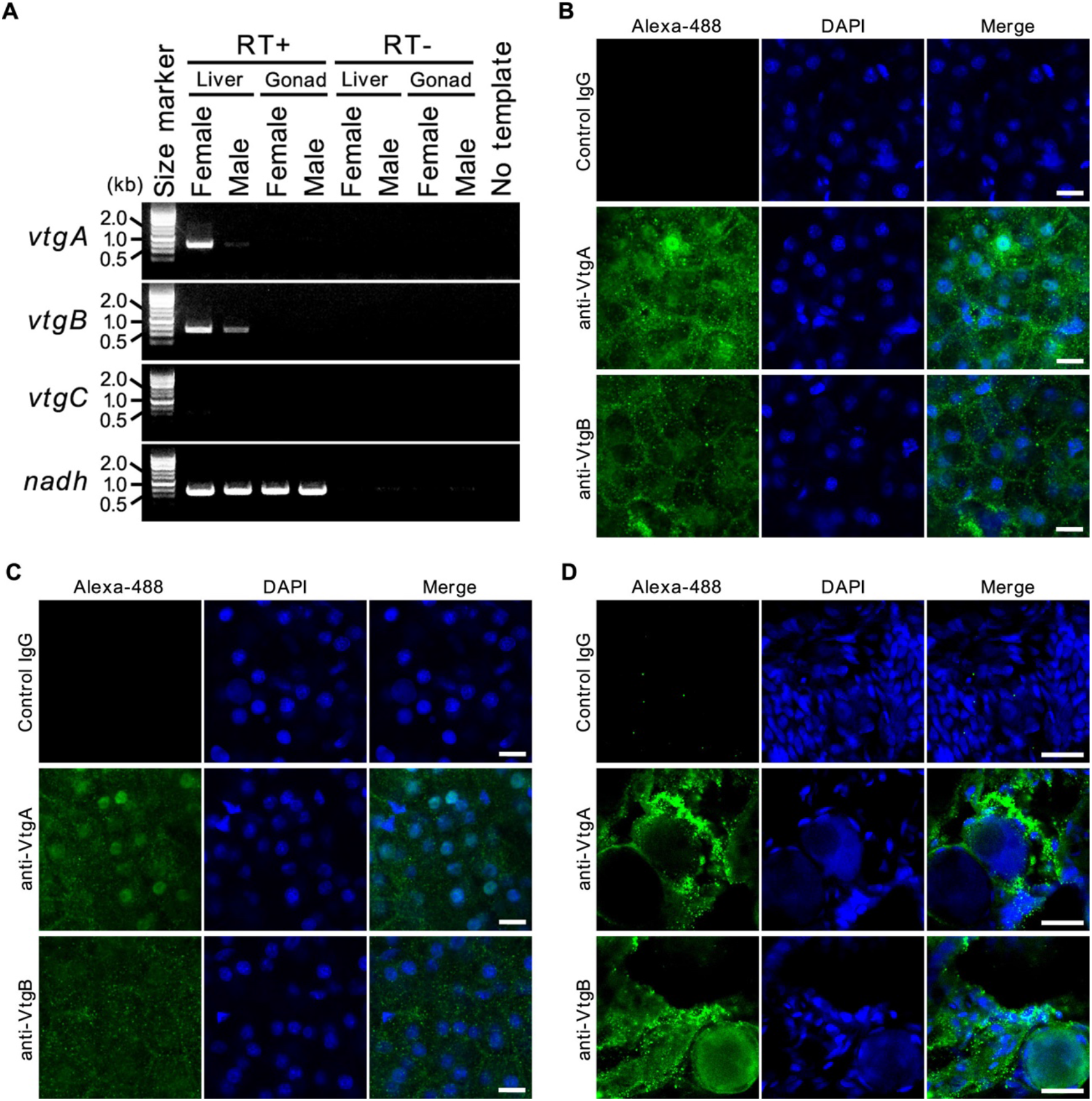
Vitellogenin synthesis and distribution in *X. eiseni*. Related to Figure 1. Additional information on *vtg* gene expression and protein distribution in female *X. eiseni* during vitellogenesis. A. RT-PCR for *vtg* genes in *X. eiseni. vtgA and B* were strongly expressed in the female liver. *vtgC* exhibited weaker expression than the other orthologous genes (see also Fig. 1B). In the male liver, expression of the *vtg* genes was weaker than that in female liver under the experimental conditions in this study. There was no *vtg* gene expression in any gonad samples. *Nicotinamide adenine dinucleotide (nadh)* was used as an internal control. RT+, reverse transcription. RT-, no reverse transcription (negative control). **B**. Fluorescence immunohistochemistry for VtgA or B in lobe edges of female liver during the vitellogenesis (non-pregnant) stage. Scale bar, 10 μm. **C**. Fluorescence immunohistochemistry for VtgA or B in lobe edges of male liver. Scale bar, 10 μm. **D**. Fluorescence immunohistochemistry for VtgA or B in the ovarian epithelium during the vitellogenesis stage. Scale bar, 20 μm.

**Figure S2.**
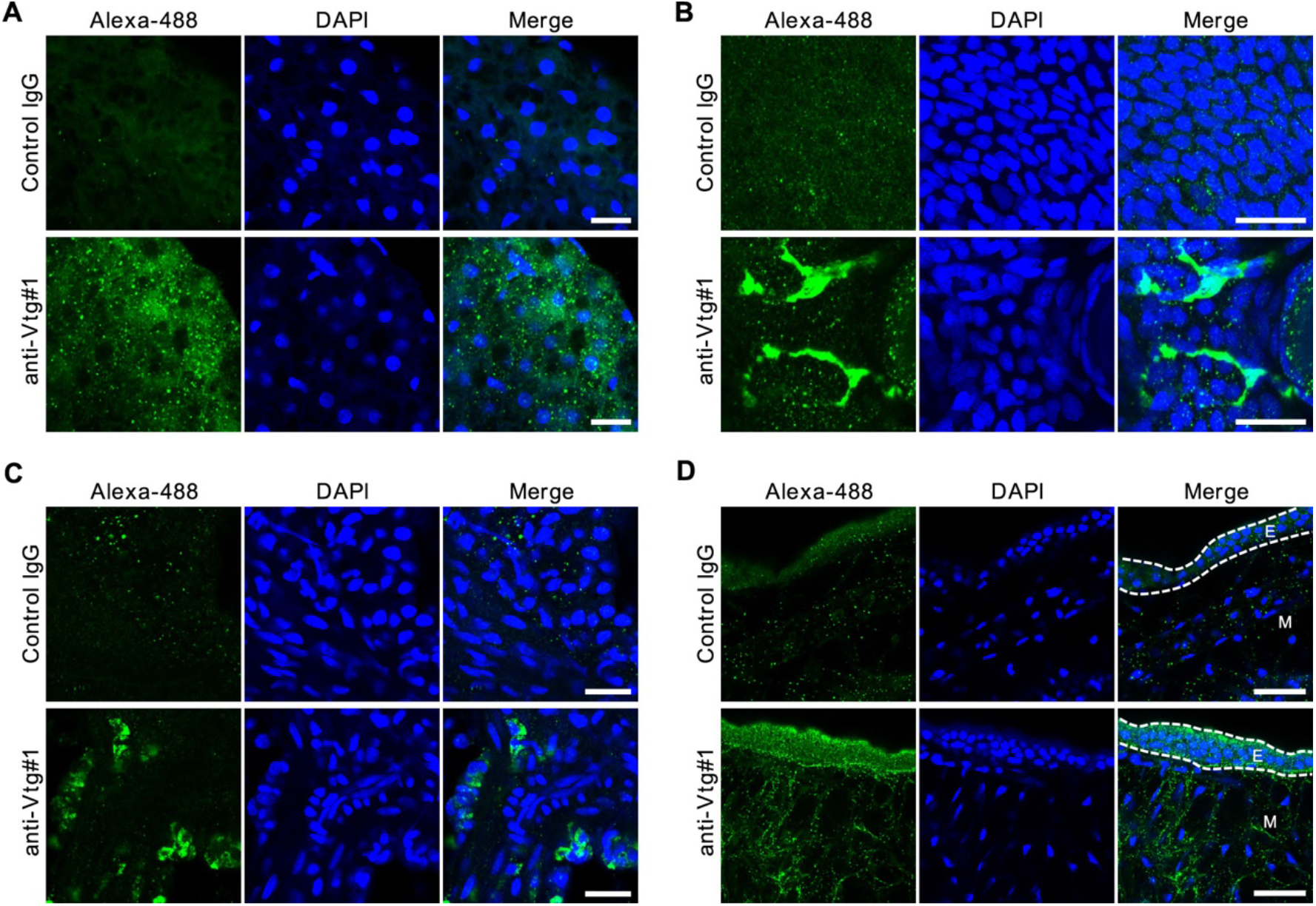
Fluorescent Immunohistochemistry using a commercial vitellogenin antibody. Related to Figures 1 and 2. Additional information on immunohistochemistry for Vtg proteins using anti-Vtg#1. **A**. Fluorescence immunohistochemistry for the female liver during the vitellogenesis stage. Scale bar, 20 μm. **B**. Fluorescence immunohistochemistry for the ovarian epithelium during the vitellogenesis stage. Scale bar, 20 μm. **C**. Fluorescence immunohistochemistry for the ovarian septum in the 3^rd^ week of gestation. Scale bar, 20 μm. **D**. Fluorescence immunohistochemistry for the trophotaenia of the intraovarian embryo in the 3^rd^ week post-fertilization. E, epidermal layer. M, mesenchyme. Scale bar, 20 μm.

**Figure S3.**
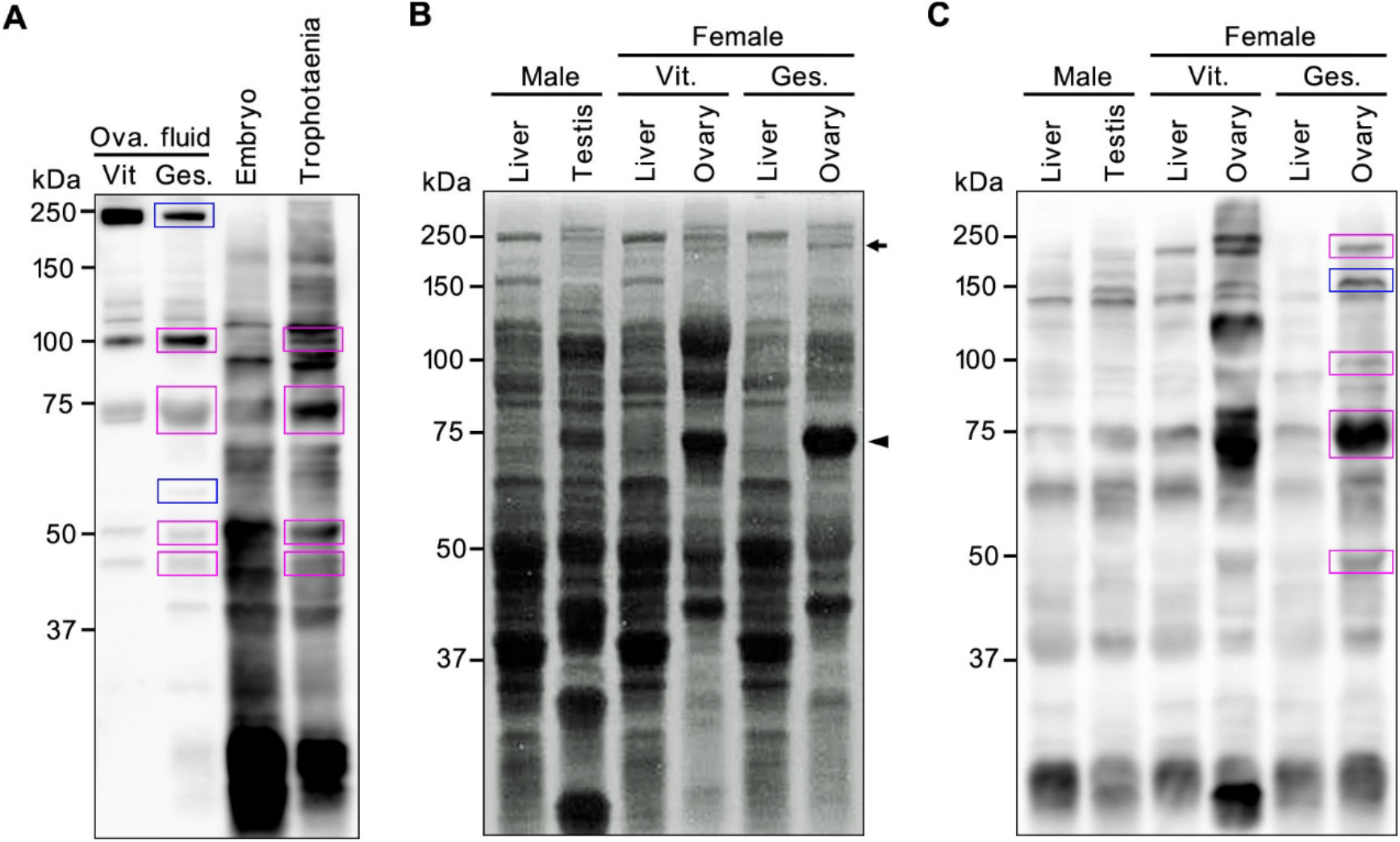
Detection of vitellogenin fragment in *X. eiseni* tissues. Related to Figure 3. Additional information on western blotting using anti-Vtg#2, and CBB staining and western blotting for Vtg in adult tissues of *X. eiseni*. **A**. Western blotting using anti-Vtg#2 for ovarian fluid and intraovarian embryo lysate extracted from pregnant female fish in the 3^rd^ week after mating. In the gestation sample, six specific signals (240-, 100-, 75-, 55-, 50-and 45-kDa) observed in Figure 3B were replicated by anti-Vtg#2. Magenta boxes indicate the signals against Vtg detected in the both the ovarian fluid and the trophotaenia. Blue boxes indicate the signals detected only in the ovarian fluid. Ova., ovarian. Vit., vitellogenesis. Ges., gestation. **B**. Electrophoresis and CBB staining of adult liver and gonad lysate extracted from male, female, and pregnant female fish in the 3^rd^ week after mating. The ovary included two major signals (240-and 75-kDa) detected in the ovarian fluid (arrow and arrowhead, see also Fig. 3A). **C**. Western blotting using anti-Vtg#2 for adult liver and gonad lysate extracted from male, female, and pregnant female fish in the 3^rd^ week after mating. The pregnant ovary included four of six signals (240-, 100-, 75-and 50-kDa) observed in the ovarian fluid (magenta boxes, see also Fig. 3B). The blue square indicates a 150-kDa signal that was not observed in the ovarian fluid or trophotaenia.

**Figure S4.**
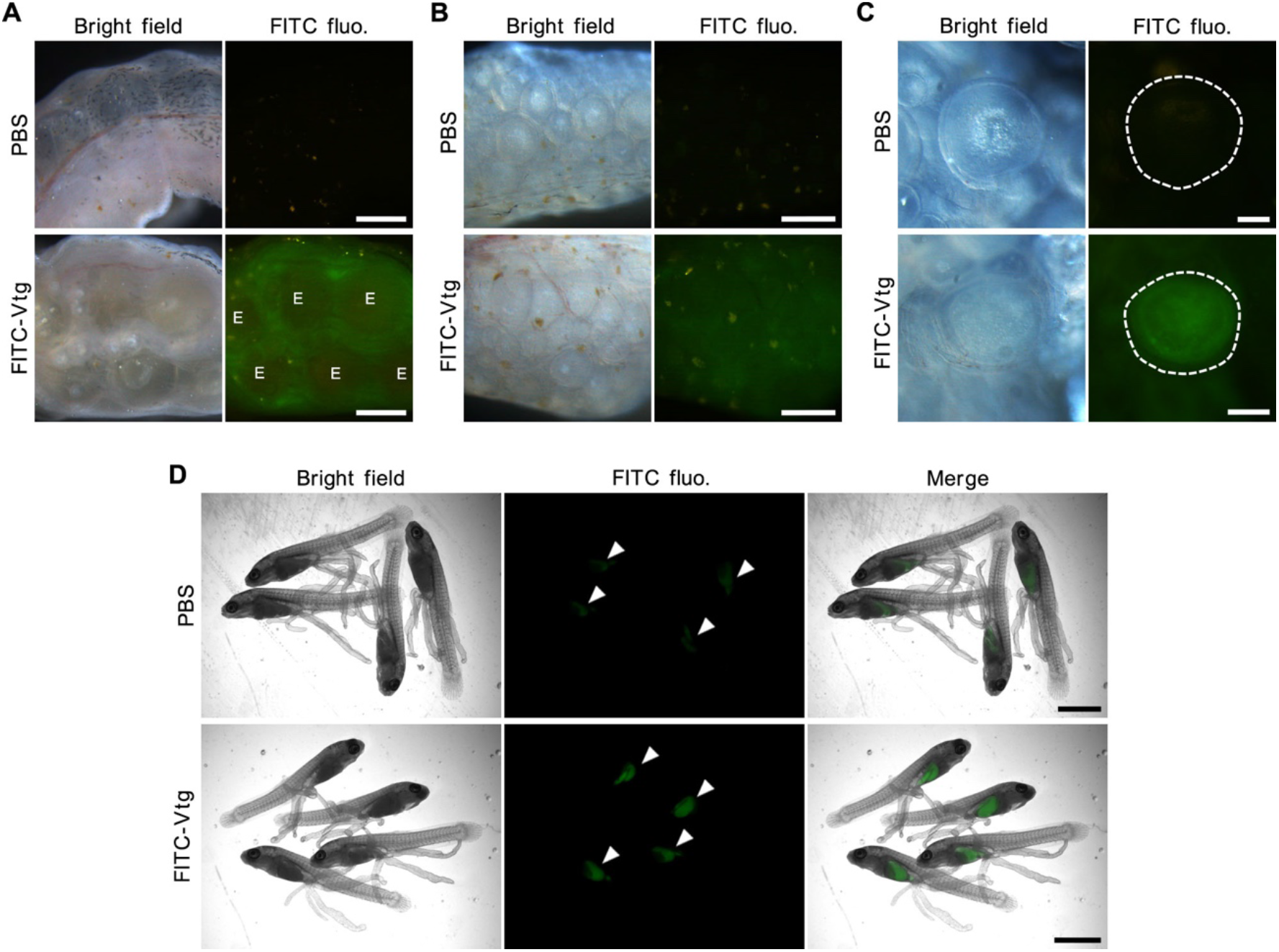
Tracing for vitellogenin transfer from abdominal cavity to ovary or intraovarian embryo. Related to Figure 4. Additional information on the Vtg tracing analysis using FITC-Vtg. **A**. Fluorescent microscopy for the ovaries extracted from female fish at 18 hours post-injection. Distinct green fluorescence was observed inside the ovary but not in the eggs extracted from the FITC-Vtg injected sample. E, egg. Scale bar, 1 mm. **B**. Fluorescent microscopy for the ovaries extracted from female fish at 90 hours post-injection. Green fluorescence was evenly observed in the ovary extracted from the FITC-Vtg injected sample. Scale bar, 500 μm. **C**. Fluorescent microscopy for the developing oocyte at 90 hours post-injection. Green fluorescence was observed in the premature yolk extracted from the FITC-Vtg injected sample. Dashed circles indicate shapes of the oocytes. Scale bar, 200 μm. **D**. Fluorescent microscopy for the intraovarian embryo at 42 hours post-injection. Green fluorescence was accumulated in the digestive tract (arrowheads) of the embryo extracted from the FITC-Vtg injected pregnant female. Scale bar, 2 mm.

**Figure S5.**
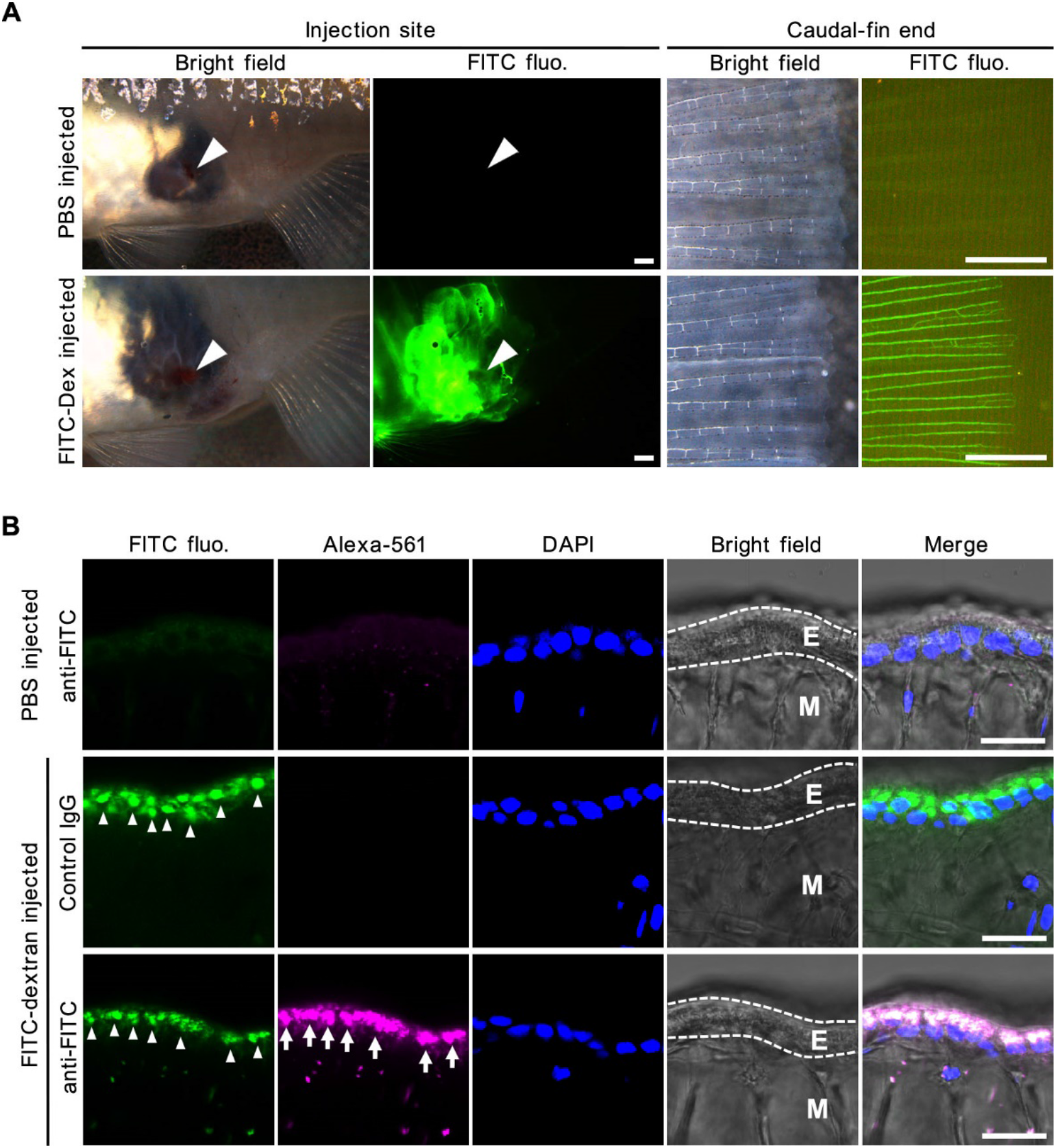
Tracing for mother-to-embryo dextran transfer in live fish. Related to Figure 4. Additional information on the macromolecule tracing analysis using FITC-dextran (MW 25,000). **A**. Fluorescent microscopy for the injected fluorescent solution in the female fish. The solutions were injected into the abdominal cavity from the gravid spot of pregnant female fish (arrowhead). The blood vessels in the caudal fin were visualized by green fluorescence of the FITC-dextran. Scale bar, 1 mm. **B**. Fluorescent microscopy of the intraovarian embryo. Green fluorescence was detected in the epidermal cell layer of the trophotaenia in the extracted embryo (arrowhead). The green fluorescence was merged to signals for anti-FITC (arrow). E, epidermal layer. M, mesenchyme. Scale bar, 10 μm.Table S1. Information for the reagents and resources used in this study.

**Table S1:**
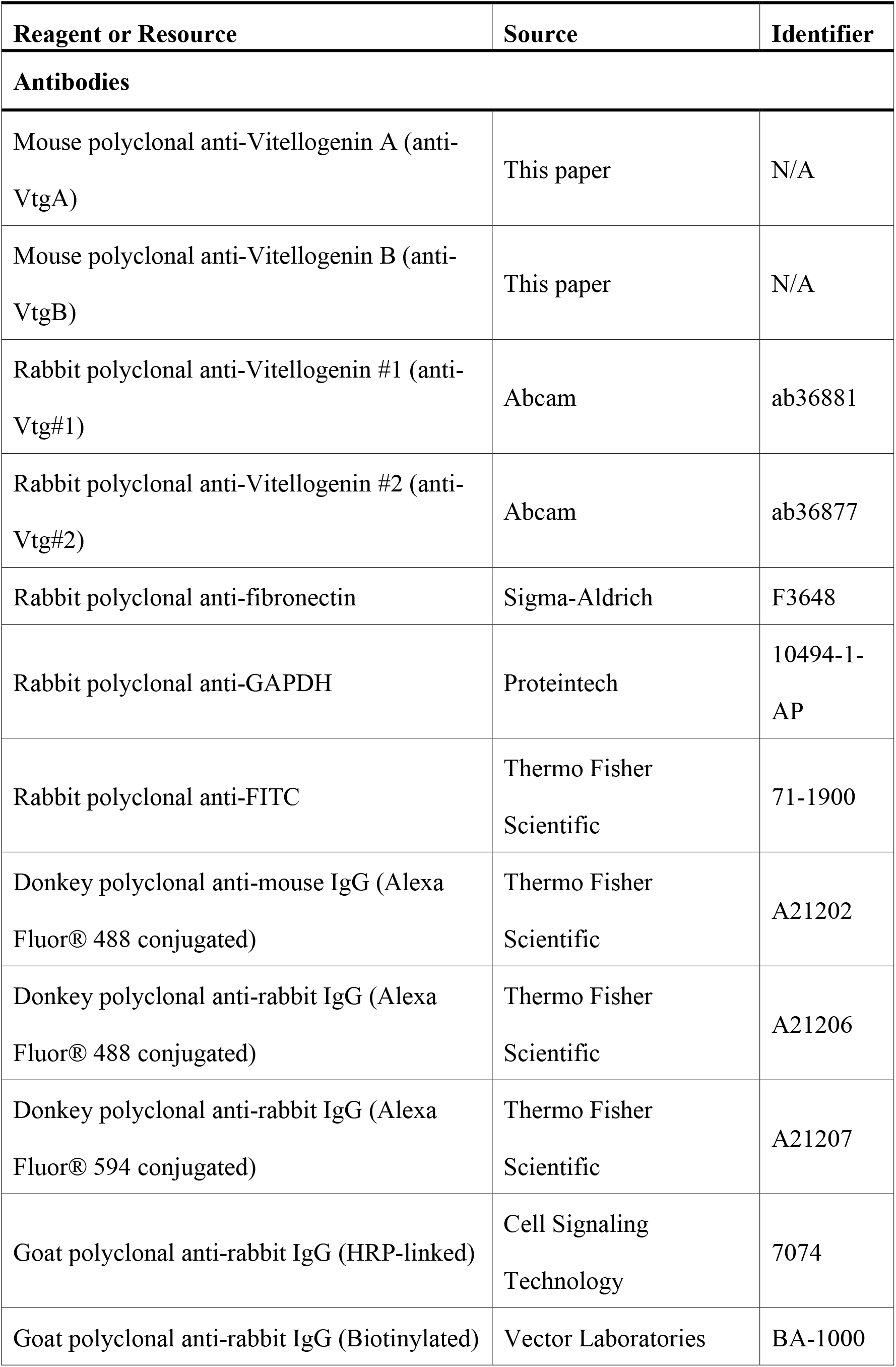
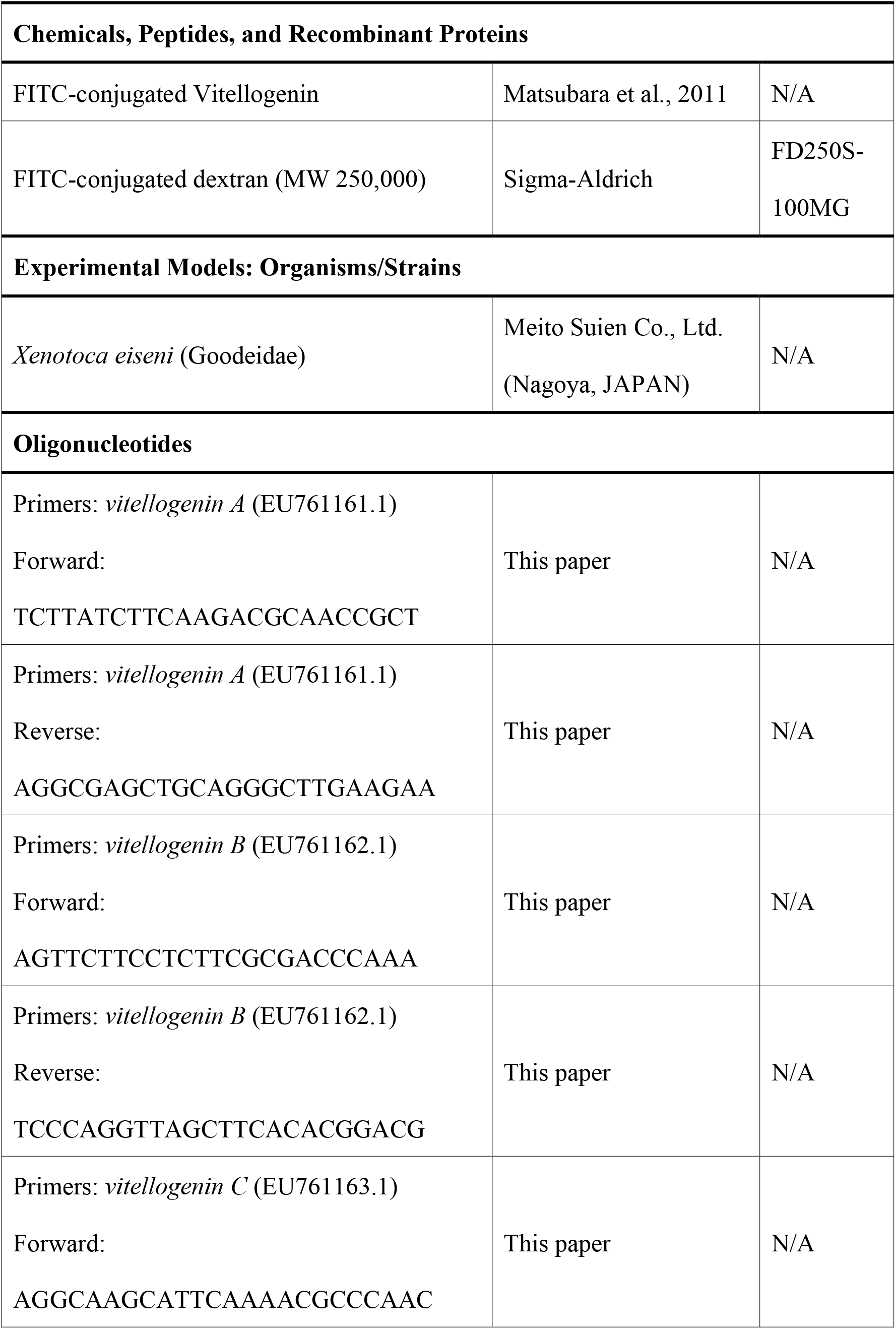
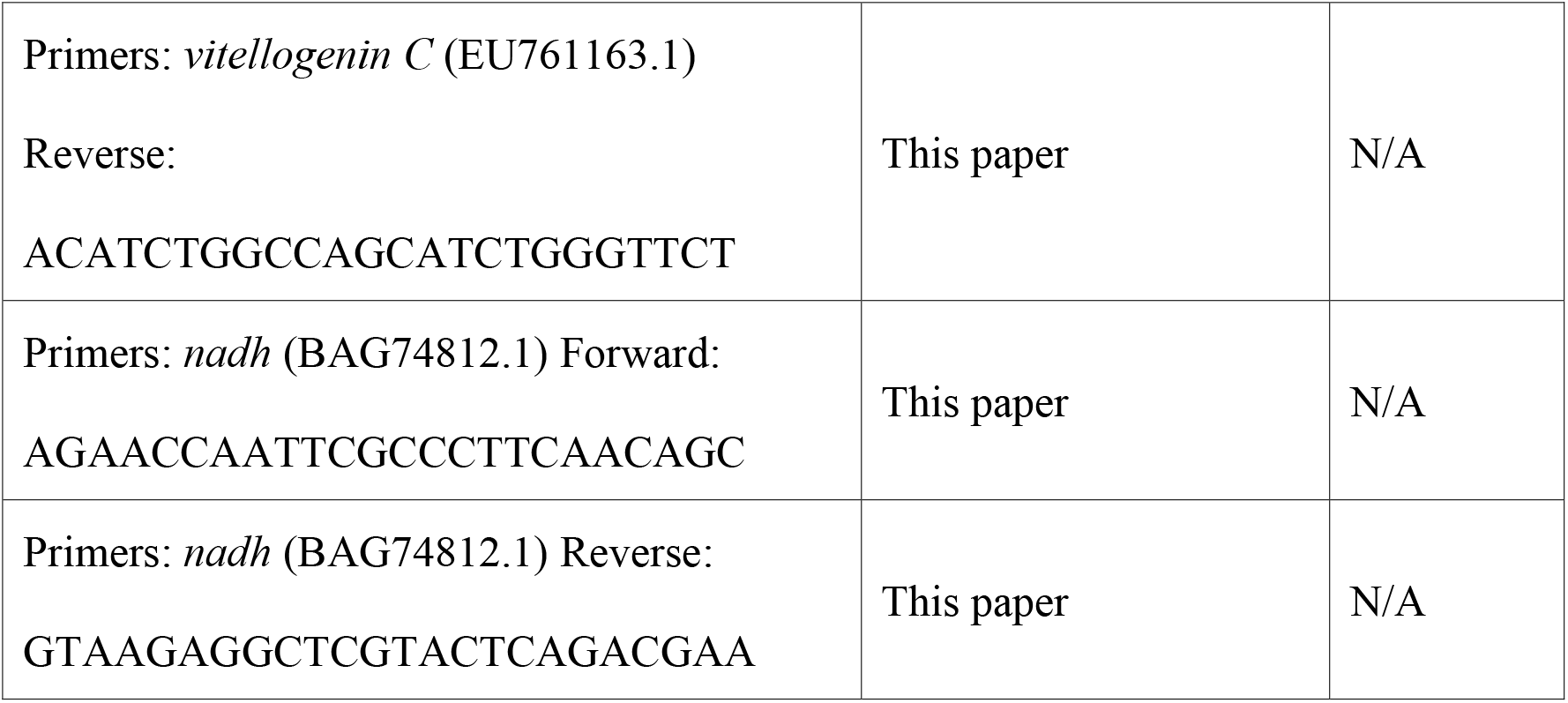
Information for the reagents and resources used in this study.

